# *In vitro* evolution of herpes simplex virus 1 (HSV-1) reveals selection for syncytia and other minor variants in Vero cell culture

**DOI:** 10.1101/760629

**Authors:** Chad V. Kuny, Christopher D. Bowen, Daniel W. Renner, Christine M. Johnston, Moriah L. Szpara

**Affiliations:** Department of Biochemistry and Molecular Biology, The Huck Institutes of the Life Sciences, and Center for Infectious Disease Dynamics, Pennsylvania State University, University Park, PA; Department of Medicine, University of Washington, Seattle, Washington, Vaccine and Infectious Disease Division, Fred Hutchinson Cancer Research Center, Seattle, WA

**Keywords:** HSV-1, evolution, minor variant, viral fitness, syncytia

## Abstract

The large dsDNA virus HSV-1 is often considered to be genetically stable, however it is known to rapidly evolve in response to strong selective pressures such as antiviral drug treatment. Deep sequencing analysis has revealed that clinical and laboratory isolates of this virus exist as populations that contain a mixture of minor alleles or variants, similar to many RNA viruses. Classical virology methods often used plaque-purified virus populations to demonstrate consistent genetic inheritance of viral traits. Plaque purification represents a severe genetic bottleneck which may or may not be representative of natural transmission of HSV-1. Since HSV-1 has a low error rate polymerase but exhibits substantial genetic diversity, the virus likely uses other mechanisms to generate genetic diversity, including recombination, contraction and expansion of tandem repeats, and imprecise DNA repair mechanisms. We sought to study the evolution of HSV-1 *in vitro*, to examine the impact of this genetic diversity in evolution, in the setting of standard laboratory conditions for viral cell culture, and in the absence of strong selective pressures. We found that a mixed population of HSV-1 was more able to evolve and adapt in culture than a plaque-purified population, though this adaptation generally occurred in a minority of the viral population. We found that certain genetic variants appeared to be positively selected for rapid growth and spread in Vero cell culture, a phenotype which was also observed in clinical samples during their first passages in culture. In the case of a minor variant that induces a visually observable syncytial phenotype, we found that changes in minor variant frequency can have a large effect on the overall phenotype of a viral population.

**Author Summary:** Herpes simplex virus type 1 (HSV-1) is a common virus, affecting over half of the adult human population, although it presents variable levels of disease burden and frequency of symptomatic recurrence. Antiviral treatments for HSV-1 infections are available, but thus far attempts at vaccine development have been foiled by insufficient immunity and/or viral escape. As a virus with a double-stranded DNA genome, HSV-1 is generally considered to be genetically stable and to have limited evolutionary potential. As these two statements are in conflict, we examined the ability of HSV-1 to evolve in a standardized cell culture setting. We utilized two HSV-1 isolates in this experiment, one with multiple viral genotypes present, which is similar to the viral populations seen in clinical settings, and one with a highly clonal viral population, which is similar to those often used in laboratory settings. After multiple rounds of replication, we analyzed the sequences of each passaged population. We found that the mixed viral population changed substantially over passage, and we were able to track specific genetic variants to phenotypic traits. By comparison, evolution in the clonal virus population was more limited. These data indicate that HSV-1 is capable of evolving rapidly, and that this evolution is facilitated by diversity in the viral population.

## Introduction

HSV-1 is a widespread pathogen in the alphaherpesvirus family [1]. It is prevalent across the world, infecting an estimated 3.7 billion people, who experience a spectrum of disease outcomes from asymptomatic infections to recurring skin lesions or rarely, encephalitis [2]. HSV-1 productively replicates in epithelial cells and then latently infects neuronal cells, leading to lifelong persistence in the host. The virus can periodically reactivate from latency to productively replicate in epithelial cells, with the potential for multiple cycles of latency and reactivation from the neuronal reservoir over the life of an infected individual [1]. This replication strategy has great advantages for viral persistence, and it also provides multiple opportunities for the virus to evolve within a host. During each active replication event, the virus may undergo selection in response to the cellular and immunological selective pressures that it faces [3–7]. Following productive replication, the progeny viruses can either be transmitted to a new host or may re-enter the nervous system of the current host, potentially expanding the latent reservoir of virus for future reactivation cycles [8,9]. As the selective pressures between epithelial and neuronal cellular environments are likely different, each cycle of latency and reactivation may create new genetic diversity in the viral population, which may be followed by a bottleneck upon the conversion to latency.

While cycles of latency and reactivation are a well-established and accepted aspect of herpesvirus biology [1,10], the concept of HSV-1 exhibiting substantial genetic diversity and evolutionary potential within an infected individual has only recently been established [6,11–15]. As with other herpesviruses, HSV-1 has a large double-stranded DNA (dsDNA) genome, and these viruses have generally been considered to be genetically stable in comparison to viruses with an RNA genome [5,16]. However an increasing body of evidence supports the idea that herpesviruses, including HSV-1, exist as a diverse genetic population *in vivo* [17–20]. A similar level of diversity may exist *in vitro* as well, depending on the method of preparation (reviewed in [14]). This genetic diversity can be generated through multiple mechanisms, including polymerase error, copy number variation, and recombination [3,4,21]. The HSV-1 polymerase has been previously demonstrated to have a low mutation rate (1 × 10^-7^ to 1 × 10^-8^ mutations per base per infectious cycle), although these studies were performed on a single gene in a unique coding region of the HSV-1 genome [3,4,22]. The HSV-1 genome consists of unique long (UL) and unique short (US) coding regions, which are flanked by large structural repeats (Internal Repeat Long/Short (IRL/S), Terminal Repeat Long/Short (TRL/S). Tandem repeats (TRs) occur frequently in the HSV-1 genome but are especially enriched in the IRL/S and TRL/S regions. Copy number variation or length fluctuations of tandem repeats and homopolymer tracts are a frequent source of genetic variation in strains of HSV-1 [23]. Repetitive regions of the HSV-1 genome also have very high G+C content, which favors recombination [21]. Recombination allows for increased genetic diversity in the absence of polymerase error, which may be especially relevant for herpesviruses [21,24,25]. These mechanisms contribute to a high level of variation in the large terminal and internal repeats, which contain genes that are critical to HSV-1 replication (ICP0, ICP4, y34.5) [1,26,27]. Crucially, HSV-1 is known to respond quite rapidly in response to strong selective pressures such as antiviral drug treatment, whether through *de novo* mutation or selection of existing variants in the population [3,28–30].

The genetic diversity of a population of HSV-1 can undergo genetic drift (e.g. Single Nucleotide Polymorphisms (SNPs), Insertions/Deletions (Indels), or TR length fluctuation) or even more dramatic genetic shifts (e.g. recombination) both *in vivo* and *in vitro*. There have been few studies that have followed a population of HSV-1 through longitudinal sampling of one person [12], or studied known transmission events [11]. These studies imply that viral population diversity can be generally maintained through transmission [11], but that the viral population can drift or adapt during subsequent rounds of latency and reactivation [12]. However, as with most viruses, HSV-1 is quite often studied in cell culture or animal models. Each of these approaches have limitations and differences from authentic replication in the natural host *in vivo*, with concomitant impacts on viral evolution and genetic diversity. As an example, HSV-1 is routinely propagated in monolayers of African green monkey kidney (Vero) cells. These cells support robust viral replication and are easily manipulated in a laboratory environment. However, these cells are not human in origin and they represent only one cell type (described as fibroblast-like) [31] of the multiple epithelial and mucosal cell types that HSV-1 encounters in a normal human infection [1]. In addition to the lack of adaptive immune control in cell culture, Vero cells lack innate immune defenses, due to a defect in interferon signaling [31–33]. Finally, HSV-1 stocks are often passaged multiple times in Vero cells to amplify the virus, without considering the potential for genetic drift from the source stock. One classic approach used by virologists to reduce the potential for genetic drift was to isolate virus from individual plaques on a monolayer, which is referred to as plaque-purification. This practice constitutes a severe genetic bottleneck, which could lead to unexpected consequences if a rare or deleterious genetic variant is unintentionally selected [26,27,34].

In this study, we utilized serial passage of two HSV-1 populations to begin to address several outstanding questions about the ability of this virus to evolve in cell culture. We utilized plaque morphology as an initial marker for genetic diversity, based on our prior observation that a frequently used HSV-1 strain (F) contained a mixture of syncytial and non-syncytial plaque phenotypes [26,27]. We also used a “purified” clonal population of the same strain of HSV-1, which was generated through plaque purification of a non-syncytial representative of strain F. By subjecting each of these viral populations to serial passage in Vero cells, we investigated the contribution of genetic diversity to viral fitness, detected the frequency and distribution within the genome of new genetic variants, and gained insight into the mechanisms underlying the generation and maintenance of genetic diversity. These results have ramifications for our understanding of the outcome of viral mutagenesis and generation of new recombinants *in vitro*, as well as shedding new light on the genetic diversity of HSV-1.

## Results

### Viral Phenotypes can Evolve over Sequential Passage in Culture

While viruses with a DNA genome are often considered genetically stable [5,16], they are able to evolve rapidly in response to strong selective pressures, such as antiviral drug selection [28–30]. It is becoming increasingly evident that these viruses, including HSV-1, can exist as populations that contain significant genetic diversity [14,17]. We sought to determine the effects of this genetic diversity on the *in vitro* evolution of HSV-1, in the context of the selective pressures present in Vero cell culture. To conduct these experiments, we used two related starting populations of HSV-1 strain F. The F-Mixed population (subsequently “Mixed”) is representative of the original HSV-1 F strain that was previously described to contain approximately 3% syncytial plaques, as well as an unexpected UL13 frameshift mutation (**Figure 1**) [26,27]. This variability in plaque phenotype was an early indication that the Mixed population was genetically heterogeneous. The F-Purified population (subsequently “Purified”) was derived from the Mixed population by three rounds of sequential plaque purification of a non-syncytial variant [26,27]. It displays a homogenous, non-syncytial plaque morphology (**Figure 1**). Each viral population was used to infect a separate monolayer of Vero cells at a MOI of 0.01. Following lysis of the cell monolayer, the resulting sample was harvested, titered, and examined for plaque morphology. This procedure is referred to as a passage for the purposes of these experiments. The titered viral stock from the first passage was used to infect the next monolayer of Vero cells at the same low MOI, and this process was repeated for 10 sequential passages. The viral population at each passage was also deep-sequenced (**Figure 1**). Low MOI conditions were chosen to allow for multiple rounds of replication in each passage, enabling the most fit viral genotypes to expand in the population.

**Figure 1.**
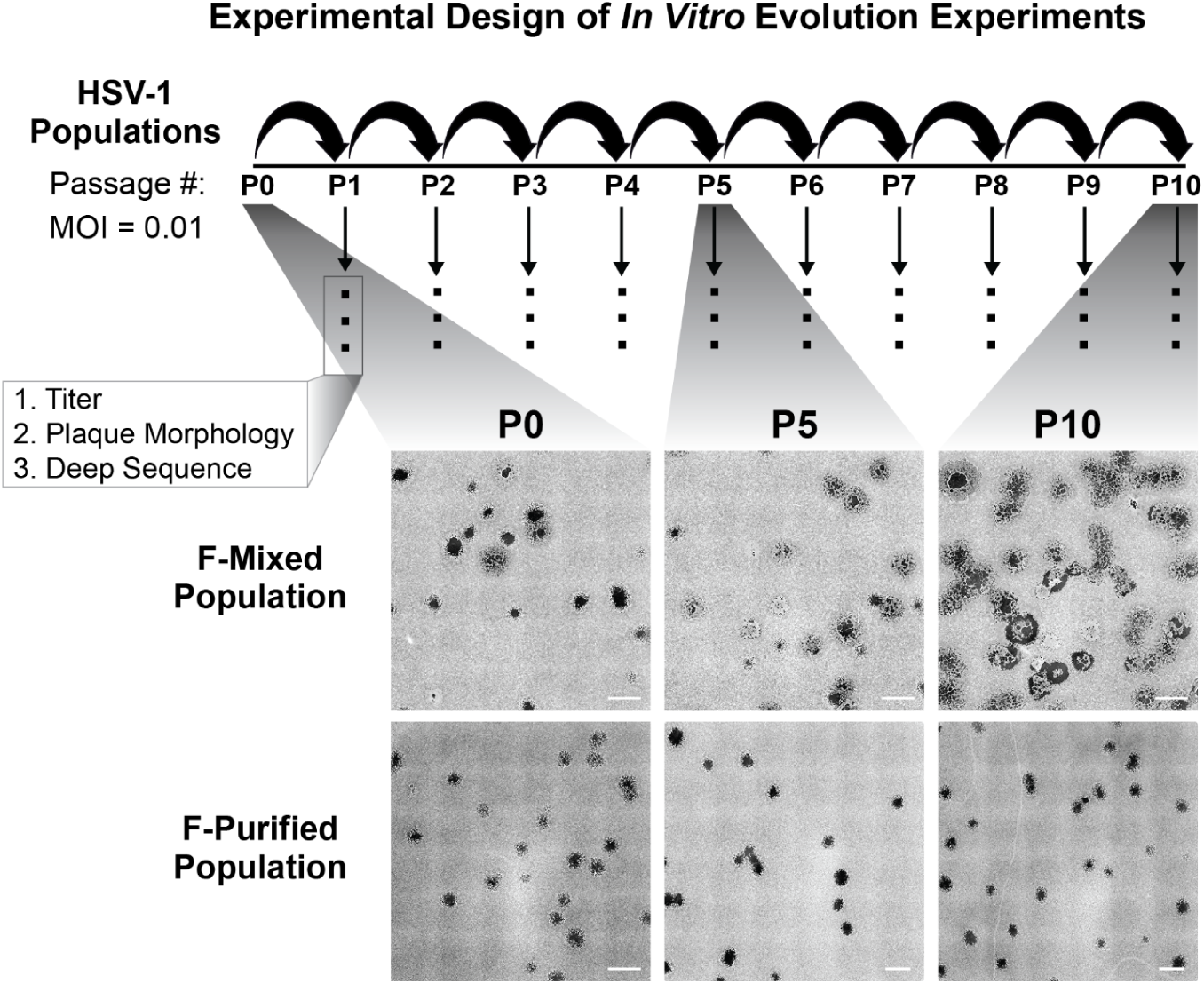
Schematic of the *In Vitro* Evolution Experiment Approach and Observed Changes in Plaque Morphology: Two populations of HSV-1 strain F were used to infect Vero cells at a multiplicity of infection (MOI) of 0.01. After infection was allowed to proceed for 72 hours, progeny viral populations were harvested. This infectious cycle is referred to as a passage, and each viral population was carried through ten passages. Following each passage, each viral population was titered, examined for plaque morphology, and prepared for sequencing. The entire series of 10 passages was performed in triplicate for the Mixed population and singly for the Purified population. All replicates were titered and quantified for plaque morphology. Deep sequencing was performed on one lineage for each starting viral population. Plaque images were obtained by plating virus at limiting dilution on monolayers of Vero cells, then fixing and staining with methylene blue at 72 hpi. Tiled images (6 x 6) were exported from Nikon NIS-Elements software, and contrast inverted using Adobe Photoshop to show plaques more clearly. Scale bars indicate 2 mm.

Over the course of ten sequential passages, we noticed dramatic changes in plaque morphology in the Mixed population (**Figure 1, Figure 2A**). However, we did not observe substantial changes in viral titer across the ten passages in either viral population (**Figure 2B**). At passage 0 (P0), we observed ∼3% syncytial plaques in the Mixed population. The frequency of syncytial plaque-forming viruses in the population increased steadily with each passage, until nearly 100% syncytial plaques were observed at passage 10 (P10). The conversion from predominantly non-syncytial to syncytial plaques in the population was observed in three independent lineages of the passaging experiment (**Figure 2A**). In contrast, the Purified population did not contain any syncytial plaques at P0, and the syncytial plaque phenotype was not acquired over the course of ten passages. Using plaque morphology as a visual marker of genetic diversity, these results suggested that the more genetically diverse Mixed population was better able to adapt or evolve over the course of this experiment.

**Figure 2.**
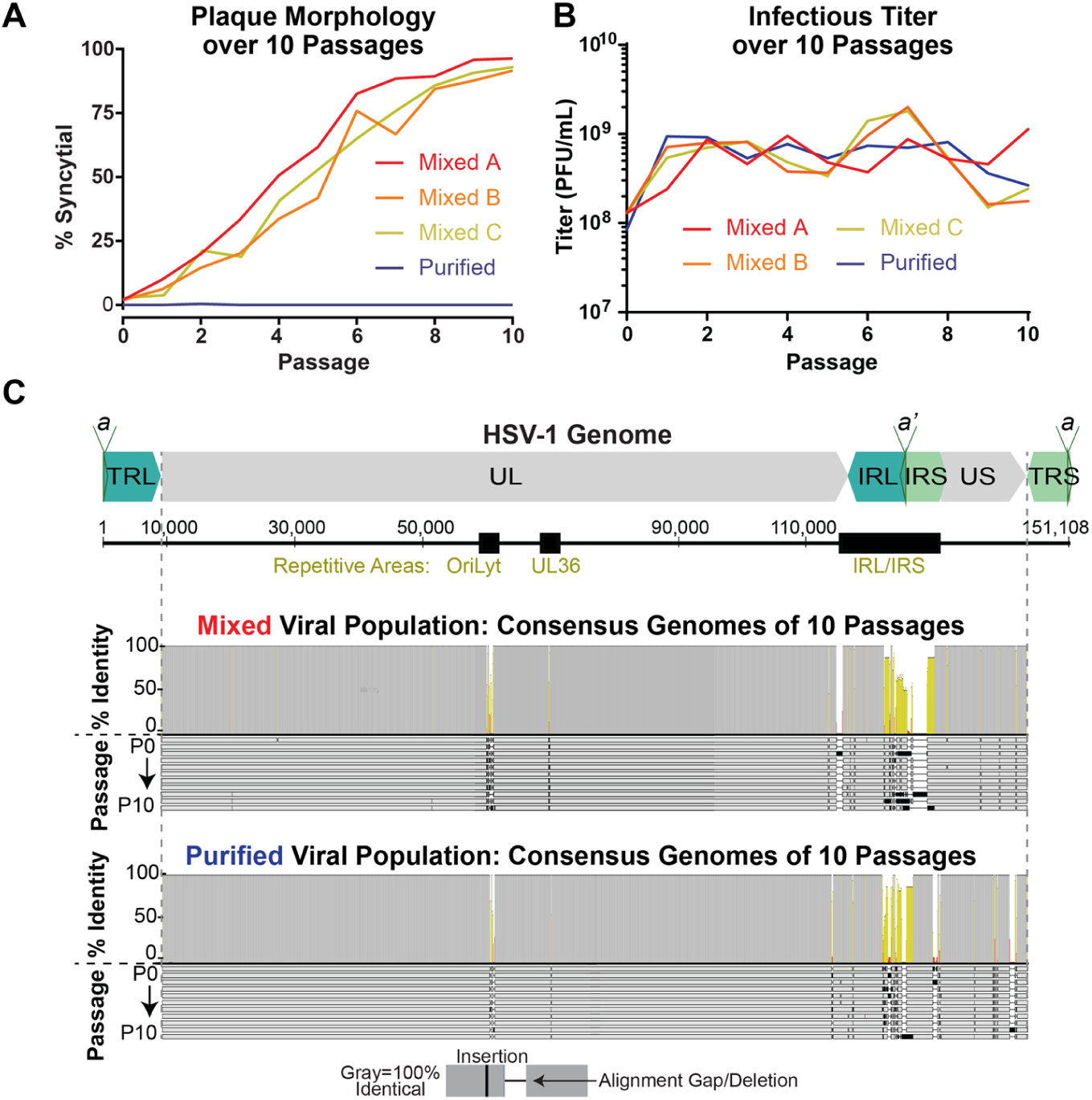
Gross Changes in Plaque Morphology Occur over Sequential Passages, while Consensus-Level Genome Changes are Limited. **A)** Plaque morphology was quantified for each passage of the Mixed and Purified viral populations. The Mixed population was passaged for three independent lineages (A, B, and C). **B)** The viral titer was measured at each passage of the Mixed and Purified lineages. **C)** A diagram of the full-length HSV-1 genome, depicts the structural repeats and major sites of tandem repeats (black boxes). An alignment of the consensus genome for each of ten passages of the Mixed and Purified viral populations are shown here. Each genome is represented by a gray bar and passages are displayed in order (P0-P10). Identical sequences are noted by gray in the identity plot above the alignment, with yellow and red indicating lesser levels of identity across the alignment. The terminal repeats (TRL/TRS) are trimmed from the consensus genomes as to not over-represent diversity in these duplicated regions. (UL, Unique Long; US, Unique Short; TRL, Terminal Repeat Long; IRL, Internal Repeat Long; IRS, Internal Repeat Short; TRS – Terminal Repeat Short, a/a’, repeat-containing site involved in genome cleavage and packaging.

### Few Changes in Consensus Genomes were Observed over 10 Passages

Because each independent Mixed lineage displayed similar changes in plaque morphology, we chose one lineage each for Mixed and Purified viral populations and used high-throughput sequencing (HTS) to examine the viral genetic diversity at each passage. Full-length, consensus genomes were assembled for each passage using a *de novo* assembly pipeline [26]. We describe these as consensus genomes because they condense the sequencing results from the entire viral population into the most common nucleotide observed at each location in the genome [14]. Therefore, these genomes represent the consensus of the sequencing reads that were aligned to each position. We found that the consensus genomes were similar overall from passage to passage within the same lineage (i.e. Mixed vs. Purified), which enabled curation and improvement of the assembled sequences. The high percent identity of a full-genome alignment derived from each set of 10 passages is presented in **Figure 2C**. While the consensus genomes were generally similar, sequence variation was observed in highly repetitive regions of these consensus genomes, where assembly of the HSV genome from short-read sequencing is technically challenging [14]. These regions include the PQ amino acid repeats in the gene encoding the tegument protein VP1/2 (UL36), the inverted repeats surrounding the lytic origin of replication (OriLyt), and the large internal and terminal copies of the repeats flanking UL and US (IRL/S and TRL/S) [27]. Surprisingly, the Mixed viral population did not display differences in genes associated with syncytia formation in the consensus genomes until passage 8 (P8). This genetic data differs from the timing of our visual detection of the syncytial plaque phenotype, which represented a majority of the observed plaques by P4. The asynchrony of consensus-level genomic data with the visually-observable phenotype motivated a deeper analysis of the sequencing results, to analyze how minor alleles in the viral population may be contributing to the observed diversity in phenotypes.

### Minor Variants Reveal Evolution in the Viral Population

Because the analysis of consensus genomes from the passaged viral populations was insufficient to explain the observed phenotypic differences, we analyzed the sequencing data for evidence of genotypes that were detected as minor alleles in the viral population. These minor variants, which we define as nucleotide alleles present in <50% of the sequencing reads at a given locus, are not otherwise represented in the consensus genomes. With sufficiently deep sequencing coverage, these variants can generally be identified with confidence. The assembled genomes across all 10 passages had an average coverage depth of 563 reads/position for the Mixed viral population and 517 reads/position for the Purified viral population. We identified nucleotide substitutions and insertions/deletions (indels) that were present in greater than 2% but less than 50% of the sequencing reads for a given sample (2% cutoff as the threshold of detection; see Methods for additional criteria.) Since the frequency of sequencing reads is correlated with the frequency of alleles in the population, this analysis revealed the genetic diversity present in each starting population, as well as the specific changes that occurred over passage, and their relative proportion in each sequential passage.

For each minor variant, we examined its position in the genome as well as its frequency in the population. We detected minor variants in each viral population, albeit at different sites and frequencies in each passage and population (**Figures 3** and **4**; see **Supplemental Files 1** and **2** for full list). Both viral populations had minor variants in highly repetitive areas of the genome, but the Mixed viral population also had multiple minor variants in the unique (e.g. UL) coding regions (**Figure 3A** and **4A**). As each virus population was passaged, the pattern of observed minor variants became more distinct. The Mixed population became more diverse, with new minor variants detected throughout the genome, and in a substantial percentage of the overall virus population (**Figure 4A**). In contrast, the vast majority of detected minor variants in the Purified population were in the highly repetitive regions of the HSV-1 genome [14] (**Figure 3B** and **4B**). These results indicated that the initially higher level of genetic diversity in the Mixed population (i.e. Mixed P0 in **Figure 3A** vs Purified P0 in **Figure 3B**) likely seeded the future evolution of that viral population.

**Figure 3.**
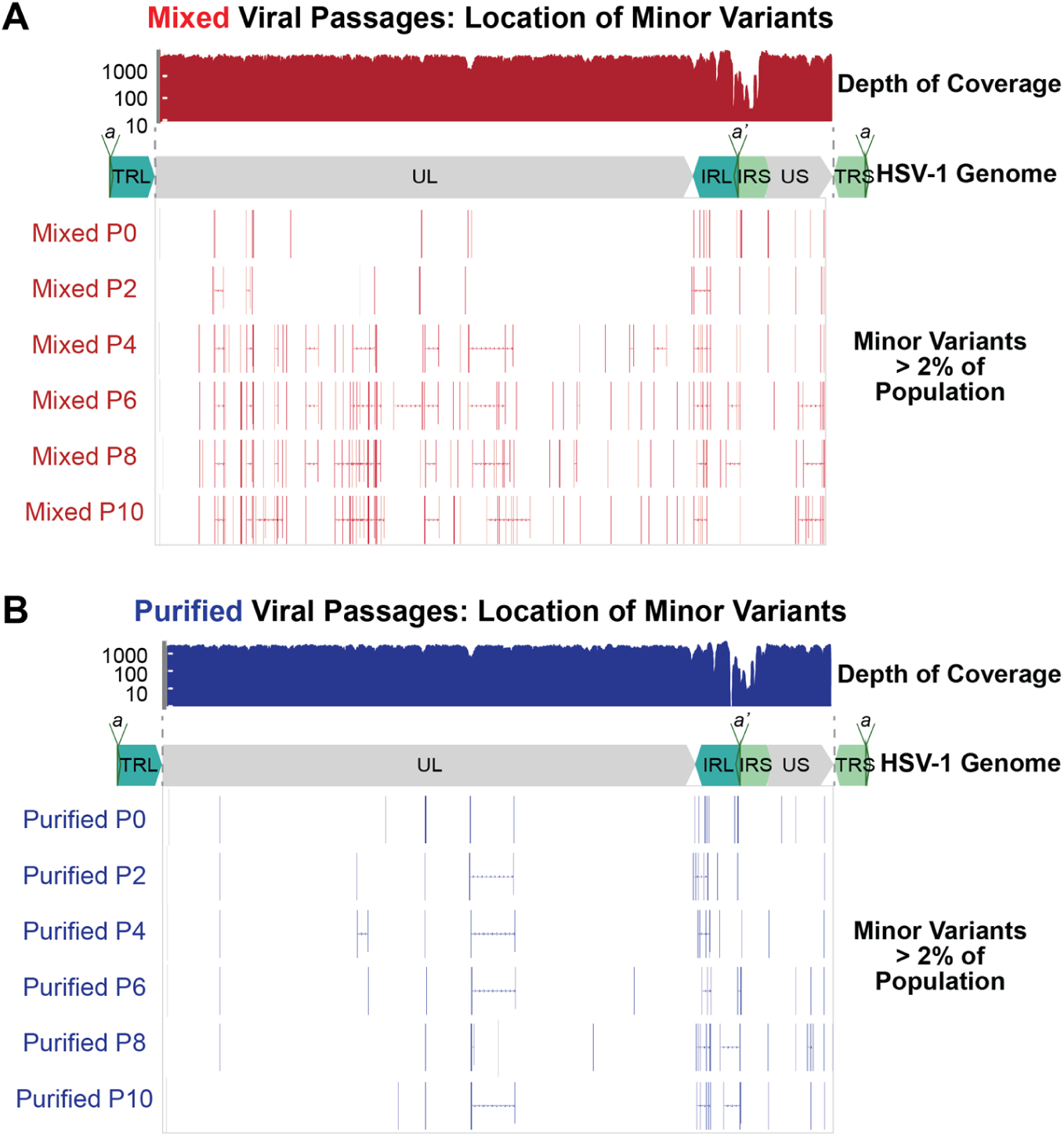
Minor Variants are Widespread in the Mixed, but not the Purified, Viral Populations. Each passage of the **A)** Mixed and **B)** Purified viral populations were sequenced and analyzed for minor variants. A subset of passages are shown here for space considerations. Each minor variant is shown as a vertical bar, and multiple variants within one gene are connected with a horizontal line. A plot of sequence read depth across the HSV-1 genome is shown for P0 of each virus population (results are representative of all passages). Across all passages, the average coverage depth across the genome was 563 reads/position for the Mixed viral populations and 517 reads/position for the Purified viral populations. A diagram of the HSV-1 genome is shown in each panel for spatial orientation.

**Figure 4.**
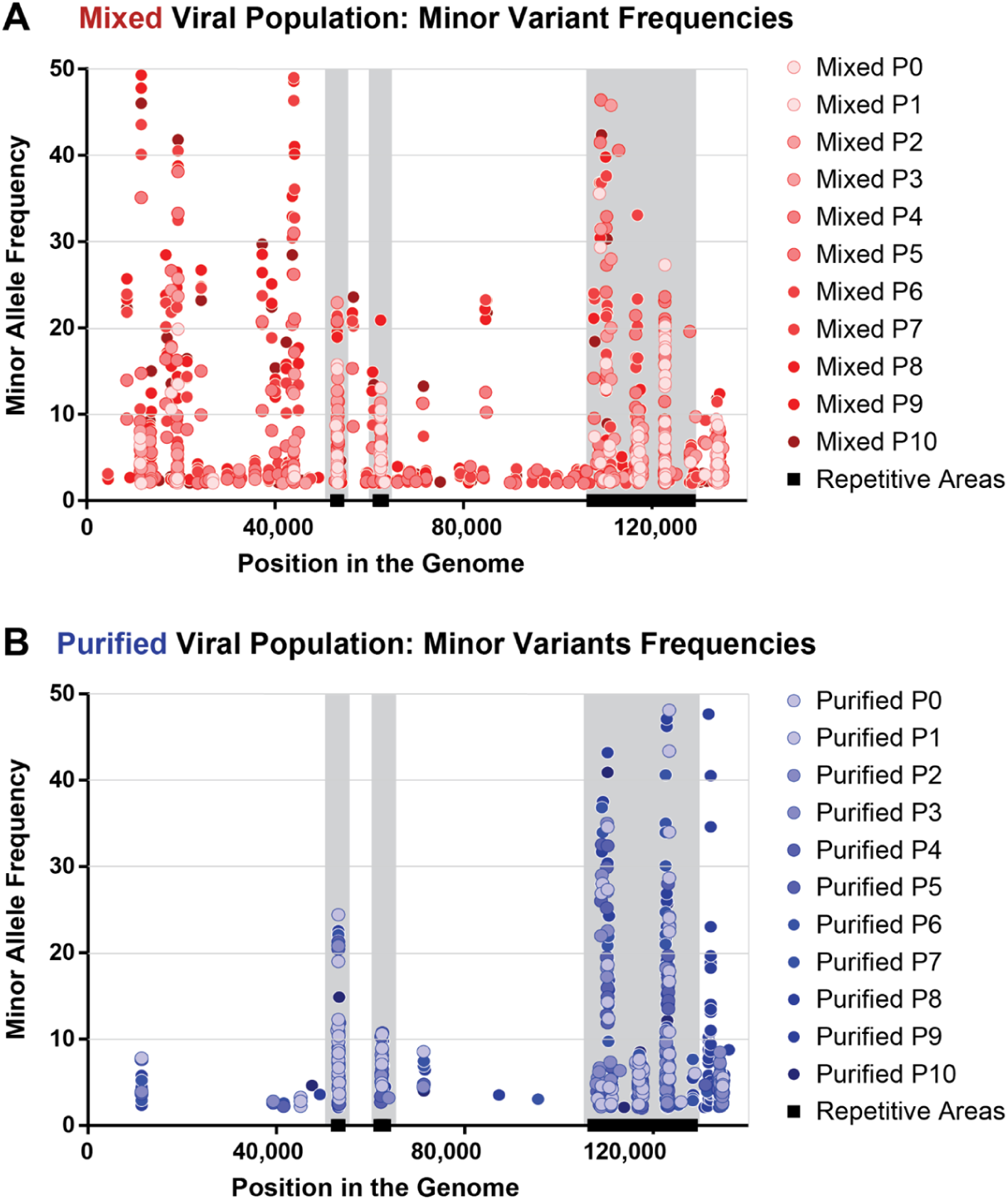
Minor Variants in the Mixed Viral Populations Occur at More Diverse Locations and at Higher Frequencies than in the Purified Viral Populations. Minor variants were plotted according to their location in the genome (x-axis), as well as by their observed frequency in each passage of these viral populations (y-axis). Variants that occurred in the same genomic location are seen as vertical columns of dots. Lighter colors indicate variants observed in earlier passages, whereas darker colors indicate later passages. Overall, a larger number and higher frequency of minor variants are observed in **A)** the Mixed viral population than in **B)** the Purified viral population. Highly repetitive regions of the HSV-1 genome are denoted by black bars (OriLyt, UL36 PQ repeats, and the internal repeat region), as in **Figure 2C**.

While the repetitive regions of herpesvirus genomes have previously been demonstrated to be areas of high variability [21,23,26,27], it is difficult to measure length fluctuations and other variations using short-read HTS technology in these regions [14]. To surmount this challenge, we used the sequential nature of these passages to identify minor variants with high confidence, due to their appearance in multiple sequential passages. We identified a number of these high-confidence minor variants (**Table 1**) in the Mixed population lineage. In contrast, we were unable to identify any such high-confidence variants of this type in the Purified population lineage. While we cannot disregard the minor variants that sporadically appear in both of these viral populations, the lack of observed consistency from passage to passage makes these sporadic minor variants more difficult to conclusively identify, especially when they occurred in the aforementioned repetitive regions. These data suggest that when HSV-1 has a more diverse starting genetic population (i.e. the Mixed population), it is more able to change in response to the pressures of a given environment.

**Table 1.**
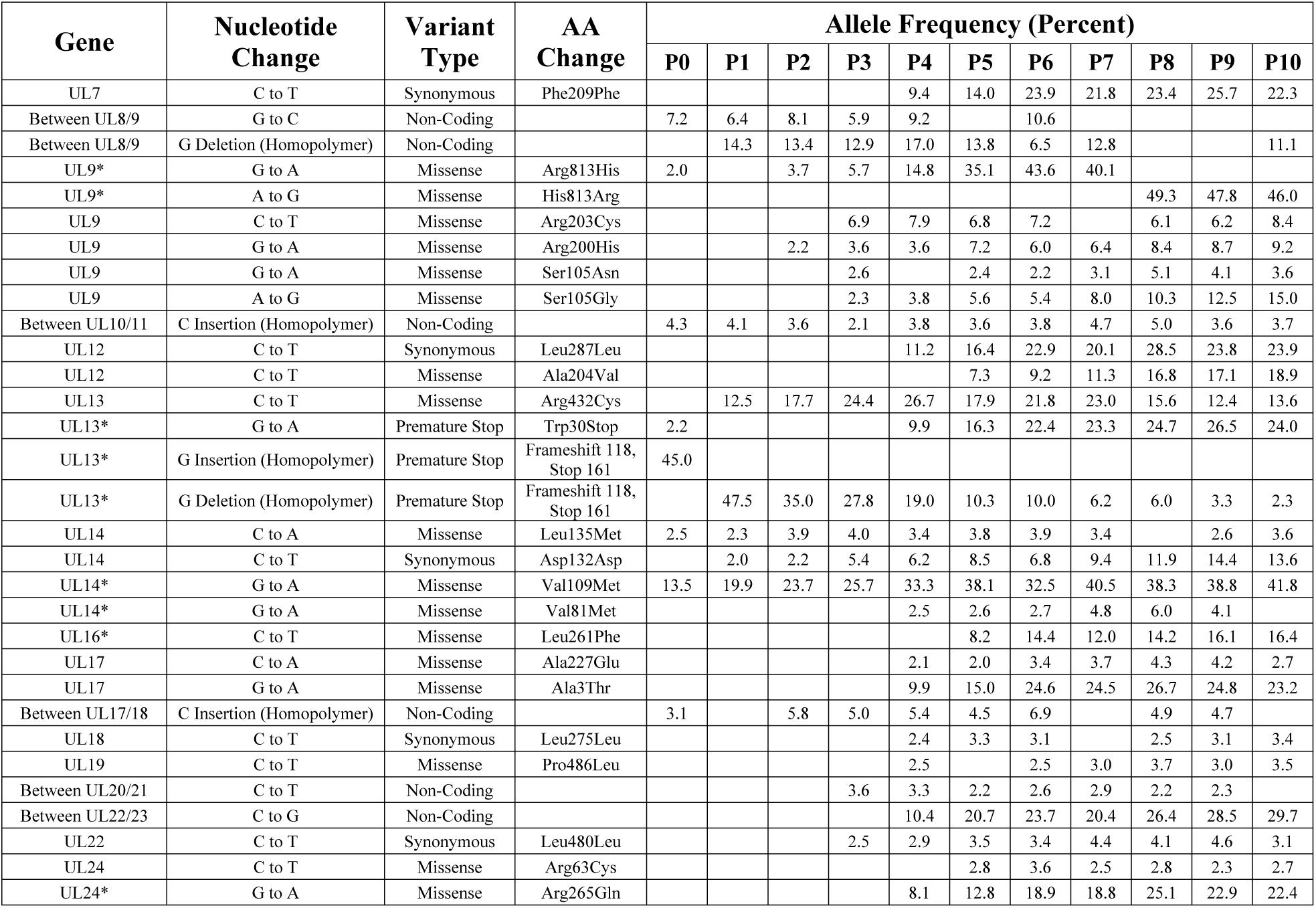
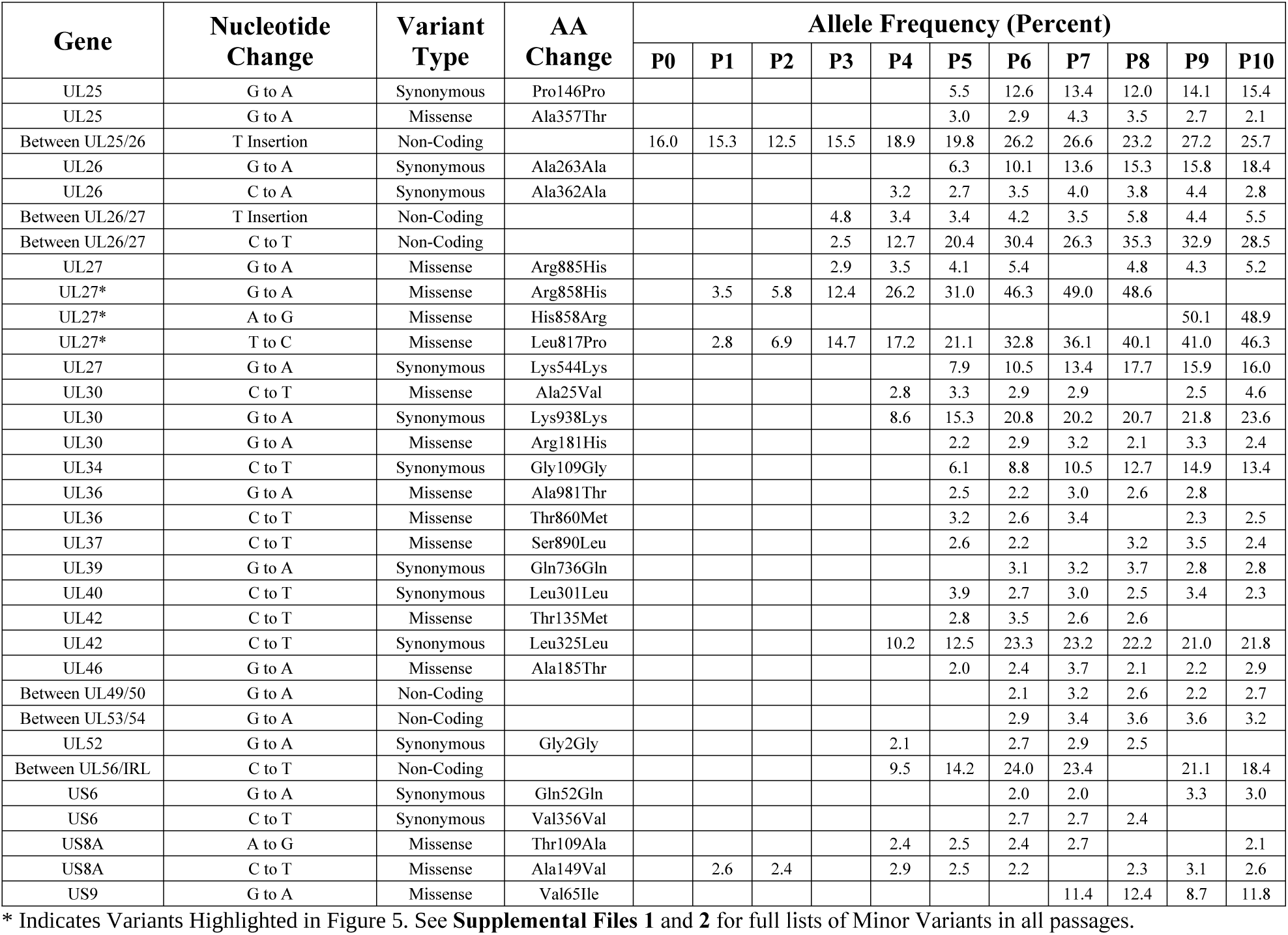
High Confidence Minor Variants in F Mixed Population. (* Indicates Variants Highlighted in Figure 5)

| Gene | Nucleotide Change | Variant Type | AA Change | Allele Frequency (Percent) |  |  |  |  |  |  |  |  |  |  |
| --- | --- | --- | --- | --- | --- | --- | --- | --- | --- | --- | --- | --- | --- | --- |
|  |  |  |  | P0 | P1 | P2 | P3 | P4 | P5 | P6 | P7 | P8 | P9 | P10 |
| UL7 | C to T | Synonymous | Phe209Phe |  |  |  |  | 9.4 | 14.0 | 23.9 | 21.8 | 23.4 | 25.7 | 22.3 |
| Between UL8/9 | G to C | Non-Coding |  | 7.2 | 6.4 | 8.1 | 5.9 | 9.2 |  | 10.6 |  |  |  |  |
| Between UL8/9 | G Deletion (Homopolymer) | Non-Coding |  |  | 14.3 | 13.4 | 12.9 | 17.0 | 13.8 | 6.5 | 12.8 |  |  | 11.1 |
| UL9* | G to A | Missense | Arg813His | 2.0 |  | 3.7 | 5.7 | 14.8 | 35.1 | 43.6 | 40.1 |  |  |  |
| UL9* | A to G | Missense | His813Arg |  |  |  |  |  |  |  |  | 49.3 | 47.8 | 46.0 |
| UL9 | C to T | Missense | Arg203Cys |  |  |  | 6.9 | 7.9 | 6.8 | 7.2 |  | 6.1 | 6.2 | 8.4 |
| UL9 | G to A | Missense | Arg200His |  |  | 2.2 | 3.6 | 3.6 | 7.2 | 6.0 | 6.4 | 8.4 | 8.7 | 9.2 |
| UL9 | G to A | Missense | Ser105Asn |  |  |  | 2.6 |  | 2.4 | 2.2 | 3.1 | 5.1 | 4.1 | 3.6 |
| UL9 | A to G | Missense | Ser105Gly |  |  |  | 2.3 | 3.8 | 5.6 | 5.4 | 8.0 | 10.3 | 12.5 | 15.0 |
| Between UL10/11 | C Insertion (Homopolymer) | Non-Coding |  | 4.3 | 4.1 | 3.6 | 2.1 | 3.8 | 3.6 | 3.8 | 4.7 | 5.0 | 3.6 | 3.7 |
| UL12 | C to T | Synonymous | Leu287Leu |  |  |  |  | 11.2 | 16.4 | 22.9 | 20.1 | 28.5 | 23.8 | 23.9 |
| UL12 | C to T | Missense | Ala204Val |  |  |  |  |  | 7.3 | 9.2 | 11.3 | 16.8 | 17.1 | 18.9 |
| UL13 | C to T | Missense | Arg432Cys |  | 12.5 | 17.7 | 24.4 | 26.7 | 17.9 | 21.8 | 23.0 | 15.6 | 12.4 | 13.6 |
| UL13* | G to A | Premature Stop | Trp30Stop | 2.2 |  |  |  | 9.9 | 16.3 | 22.4 | 23.3 | 24.7 | 26.5 | 24.0 |
| UL13* | G Insertion (Homopolymer) | Premature Stop | Frameshift 118, Stop 161 | 45.0 |  |  |  |  |  |  |  |  |  |  |
| UL13* | G Deletion (Homopolymer) | Premature Stop | Frameshift 118, Stop 161 |  | 47.5 | 35.0 | 27.8 | 19.0 | 10.3 | 10.0 | 6.2 | 6.0 | 3.3 | 2.3 |
| UL14 | C to A | Missense | Leu135Met | 2.5 | 2.3 | 3.9 | 4.0 | 3.4 | 3.8 | 3.9 | 3.4 |  | 2.6 | 3.6 |
| UL14 | C to T | Synonymous | Asp132Asp |  | 2.0 | 2.2 | 5.4 | 6.2 | 8.5 | 6.8 | 9.4 | 11.9 | 14.4 | 13.6 |
| UL14* | G to A | Missense | Val109Met | 13.5 | 19.9 | 23.7 | 25.7 | 33.3 | 38.1 | 32.5 | 40.5 | 38.3 | 38.8 | 41.8 |
| UL14* | G to A | Missense | Val81Met |  |  |  |  | 2.5 | 2.6 | 2.7 | 4.8 | 6.0 | 4.1 |  |
| UL16* | C to T | Missense | Leu261Phe |  |  |  |  |  | 8.2 | 14.4 | 12.0 | 14.2 | 16.1 | 16.4 |
| UL17 | C to A | Missense | Ala227Glu |  |  |  |  | 2.1 | 2.0 | 3.4 | 3.7 | 4.3 | 4.2 | 2.7 |
| UL17 | G to A | Missense | Ala3Thr |  |  |  |  | 9.9 | 15.0 | 24.6 | 24.5 | 26.7 | 24.8 | 23.2 |
| Between UL17/18 | C Insertion (Homopolymer) | Non-Coding |  | 3.1 |  | 5.8 | 5.0 | 5.4 | 4.5 | 6.9 |  | 4.9 | 4.7 |  |
| UL18 | C to T | Synonymous | Leu275Leu |  |  |  |  | 2.4 | 3.3 | 3.1 |  | 2.5 | 3.1 | 3.4 |
| UL19 | C to T | Missense | Pro486Leu |  |  |  |  | 2.5 |  | 2.5 | 3.0 | 3.7 | 3.0 | 3.5 |
| Between UL20/21 | C to T | Non-Coding |  |  |  |  | 3.6 | 3.3 | 2.2 | 2.6 | 2.9 | 2.2 | 2.3 |  |
| Between UL22/23 | C to G | Non-Coding |  |  |  |  |  | 10.4 | 20.7 | 23.7 | 20.4 | 26.4 | 28.5 | 29.7 |
| UL22 | C to T | Synonymous | Leu480Leu |  |  |  | 2.5 | 2.9 | 3.5 | 3.4 | 4.4 | 4.1 | 4.6 | 3.1 |
| UL24 | C to T | Missense | Arg63Cys |  |  |  |  |  | 2.8 | 3.6 | 2.5 | 2.8 | 2.3 | 2.7 |
| UL24* | G to A | Missense | Arg265Gln |  |  |  |  | 8.1 | 12.8 | 18.9 | 18.8 | 25.1 | 22.9 | 22.4 |
|  |  |  |  | P0 | P1 | P2 | P3 | P4 | P5 | P6 | P7 | P8 | P9 | P10 |
| UL25 | G to A | Synonymous | Pro146Pro |  |  |  |  |  | 5.5 | 12.6 | 13.4 | 12.0 | 14.1 | 15.4 |
| UL25 | G to A | Missense | Ala357Thr |  |  |  |  |  | 3.0 | 2.9 | 4.3 | 3.5 | 2.7 | 2.1 |
| Between UL25/26 | T Insertion | Non-Coding |  | 16.0 | 15.3 | 12.5 | 15.5 | 18.9 | 19.8 | 26.2 | 26.6 | 23.2 | 27.2 | 25.7 |
| UL26 | G to A | Synonymous | Ala263Ala |  |  |  |  |  | 6.3 | 10.1 | 13.6 | 15.3 | 15.8 | 18.4 |
| UL26 | C to A | Synonymous | Ala362Ala |  |  |  |  | 3.2 | 2.7 | 3.5 | 4.0 | 3.8 | 4.4 | 2.8 |
| Between UL26/27 | T Insertion | Non-Coding |  |  |  |  | 4.8 | 3.4 | 3.4 | 4.2 | 3.5 | 5.8 | 4.4 | 5.5 |
| Between UL26/27 | C to T | Non-Coding |  |  |  |  | 2.5 | 12.7 | 20.4 | 30.4 | 26.3 | 35.3 | 32.9 | 28.5 |
| UL27 | G to A | Missense | Arg885His |  |  |  | 2.9 | 3.5 | 4.1 | 5.4 |  | 4.8 | 4.3 | 5.2 |
| UL27* | G to A | Missense | Arg858His |  | 3.5 | 5.8 | 12.4 | 26.2 | 31.0 | 46.3 | 49.0 | 48.6 |  |  |
| UL27* | A to G | Missense | His858Arg |  |  |  |  |  |  |  |  |  | 50.1 | 48.9 |
| UL27* | T to C | Missense | Leu817Pro |  | 2.8 | 6.9 | 14.7 | 17.2 | 21.1 | 32.8 | 36.1 | 40.1 | 41.0 | 46.3 |
| UL27 | G to A | Synonymous | Lys544Lys |  |  |  |  |  | 7.9 | 10.5 | 13.4 | 17.7 | 15.9 | 16.0 |
| UL30 | C to T | Missense | Ala25Val |  |  |  |  | 2.8 | 3.3 | 2.9 | 2.9 |  | 2.5 | 4.6 |
| UL30 | G to A | Synonymous | Lys938Lys |  |  |  |  | 8.6 | 15.3 | 20.8 | 20.2 | 20.7 | 21.8 | 23.6 |
| UL30 | G to A | Missense | Arg181His |  |  |  |  |  | 2.2 | 2.9 | 3.2 | 2.1 | 3.3 | 2.4 |
| UL34 | C to T | Synonymous | Gly109Gly |  |  |  |  |  | 6.1 | 8.8 | 10.5 | 12.7 | 14.9 | 13.4 |
| UL36 | G to A | Missense | Ala981Thr |  |  |  |  |  | 2.5 | 2.2 | 3.0 | 2.6 | 2.8 |  |
| UL36 | C to T | Missense | Thr860Met |  |  |  |  |  | 3.2 | 2.6 | 3.4 |  | 2.3 | 2.5 |
| UL37 | C to T | Missense | Ser890Leu |  |  |  |  |  | 2.6 | 2.2 |  | 3.2 | 3.5 | 2.4 |
| UL39 | G to A | Synonymous | Gln736Gln |  |  |  |  |  |  | 3.1 | 3.2 | 3.7 | 2.8 | 2.8 |
| UL40 | C to T | Synonymous | Leu301Leu |  |  |  |  |  | 3.9 | 2.7 | 3.0 | 2.5 | 3.4 | 2.3 |
| UL42 | C to T | Missense | Thr135Met |  |  |  |  |  | 2.8 | 3.5 | 2.6 | 2.6 |  |  |
| UL42 | C to T | Synonymous | Leu325Leu |  |  |  |  | 10.2 | 12.5 | 23.3 | 23.2 | 22.2 | 21.0 | 21.8 |
| UL46 | G to A | Missense | Ala185Thr |  |  |  |  |  | 2.0 | 2.4 | 3.7 | 2.1 | 2.2 | 2.9 |
| Between UL49/50 | G to A | Non-Coding |  |  |  |  |  |  |  | 2.1 | 3.2 | 2.6 | 2.2 | 2.7 |
| Between UL53/54 | G to A | Non-Coding |  |  |  |  |  |  |  | 2.9 | 3.4 | 3.6 | 3.6 | 3.2 |
| UL52 | G to A | Synonymous | Gly2Gly |  |  |  |  | 2.1 |  | 2.7 | 2.9 | 2.5 |  |  |
| Between UL56/IRL | C to T | Non-Coding |  |  |  |  |  | 9.5 | 14.2 | 24.0 | 23.4 |  | 21.1 | 18.4 |
| US6 | G to A | Synonymous | Gln52Gln |  |  |  |  |  |  | 2.0 | 2.0 |  | 3.3 | 3.0 |
| US6 | C to T | Synonymous | Val356Val |  |  |  |  |  |  | 2.7 | 2.7 | 2.4 |  |  |
| US8A | A to G | Missense | Thr109Ala |  |  |  |  | 2.4 | 2.5 | 2.4 | 2.7 |  |  | 2.1 |
| US8A | C to T | Missense | Ala149Val |  | 2.6 | 2.4 |  | 2.9 | 2.5 | 2.2 |  | 2.3 | 3.1 | 2.6 |
| US9 | G to A | Missense | Val65Ile |  |  |  |  |  |  |  | 11.4 | 12.4 | 8.7 | 11.8 |

\* Indicates Variants Highlighted in Figure 5. See **Supplemental Files 1** and **2** for full lists of Minor Variants in all passages.

### Frequency of Minor Variants May Reveal Selective Pressures *In Vitro*

We next investigated the types of minor variants that were present and how their frequencies changed over sequential passages of the viral population. We found that most of the high-confidence minor variants were missense mutations, though examples of non-coding and synonymous or silent mutations were also identified (**Table 1**). Many minor variants increased as a percentage of the population over sequential passage, while others seemed to reach an equilibrium level of standing variation (**Figure 5A** and **Table 1**). We did not observe any minor variants that were clearly linked as a haplotype, in that their frequencies did not change in parallel. Accordingly, we treated each minor variant an individual contributor to the population. Minor variants that significantly increased in the viral population potentially conferred a selective advantage under these experimental conditions. This outcome is most clearly illustrated by the syncytia-inducing variants in UL27 (glycoprotein B, gB), which match previously described spontaneous mutations observed in this and other strains of HSV-1 [35–38].

**Figure 5.**
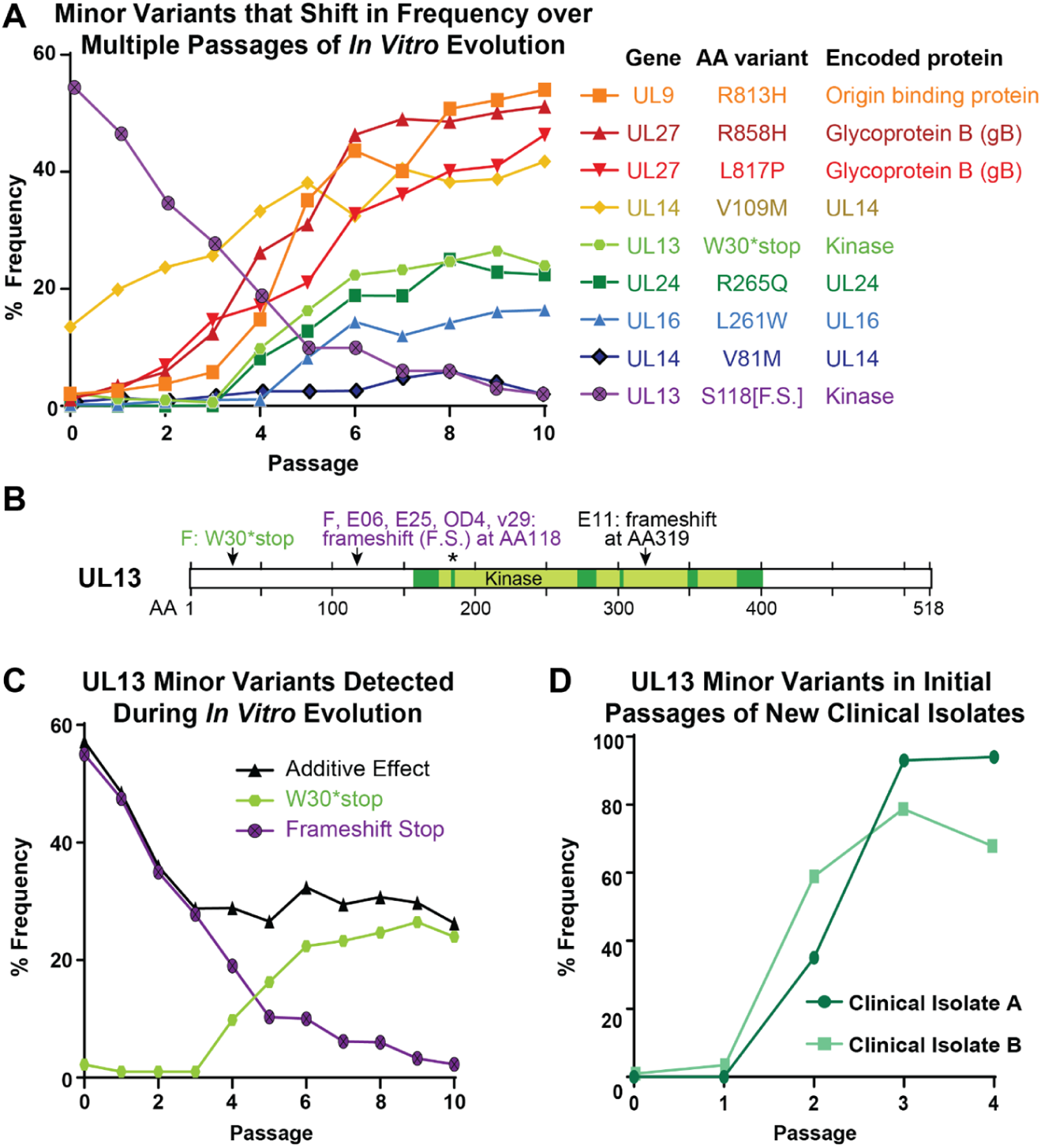
Minor Variant Dynamics Shift over Sequential Passages *In Vitro*. **A)** A subset of the observed high-confidence minor variants in the Mixed population were plotted by their frequency in the viral population over passage (see **Table 1** for full list of minor variants). Each variant would cause a change in the translated protein, whether through premature stop (UL13 S118[F.S.], W30*stop) or a missense variant (all others). Variants and their encoded proteins are listed in the legend according to their frequency at passage 10 (P10). **B)** A diagram of the UL13 protein (518 AA in length) depicts the location of the six previously described catalytic domains of this kinase [63], with an asterisk denoting the catalytic lysine where a single point mutation can disrupt kinase activity [42,43]. The minor variants observed in this study, and consensus-level mutations observed in prior studies, are indicated by arrows and labels. GenBank accessions: strain F, GU734771 [27]; E25, HM585506; E06, HM585496; E11, HM585500 [all three from [23]]; OD4, JN420342 [21]. **C)** The minor variants observed in the UL13 gene were separated for specific analysis. Each variant is predicted to produce a catalytically-inactive UL13 kinase. Additive Effect refers to the arithmetic addition of the frequency of each of these two individual variants. **D)** New clinical isolates of HSV-1 were passaged in Vero cells, genome sequenced, and examined for minor variants in the UL13 gene at each passage. The frequency of genetic variants that would produce a catalytically-inactive UL13 kinase are plotted over each passage.

Other minor variants appeared to reach an equilibrium in frequency within the viral population, which is a more difficult pattern to interpret. We found a particularly intriguing example of equilibration within the population with minor variants present in the UL13 gene. UL13 is the HSV-1 encoded representative of the Conserved Herpesvirus-encoded Protein Kinases (CHPK). UL13 kinase activity is dispensable *in vitro* [39], but it is required for axonal transport *in vivo* [40]. The kinase domain of this enzyme is located in the C-terminal end of the protein, with the crucial catalytic lysine located at amino acid 176 (**Figure 5B**) [41–43]. We identified two distinct minor variants that would cause premature termination of UL13 and would not be anticipated to have any kinase activity. We refer to these variants as W30*stop, a missense mutation, and Frameshift Stop (or S118[F.S.]), an indel in a homopolymer (**Figure 5B** and **5C**). The Frameshift Stop insertion was present in a slight majority of the initial population (51%) and it rapidly declined in frequency over sequential passages. However, the W30*stop variant increased in frequency in the population over the same passages. In combination, these two presumably defective variants of the UL13 gene added up to a stable level of ∼30% of the viral population. Thus, we observed a tolerance and/or a selection of UL13 variants that would presumably produce inactive kinases (**Figure 5B** and **5C**).

The presence of defective UL13 variants in HSV-1 populations is not unique to the F strain. In a recent publication by our group that compared a cultured clinical isolate to direct-from-patient swab-based sequencing of a genital HSV-1 lesion, we detected UL13 minor variants only in the cultured material [12]. We extended this investigation to two newly isolated clinical genital HSV-1 isolates, and likewise detected UL13-inactivating minor variants in these cultured populations. These variants rose in frequency over the initial passages *in vitro*, reaching substantially higher levels, in fewer passages, than the rate of change observed in the Mixed population (**Figure 5D**). It is as yet unknown why these viral populations would maintain standing variation in the UL13 gene, and/or favor the inactivation of UL13 in Vero cell culture.

### Syncytial Phenotype is Favored in Vero Cells

While the role of UL13 in Vero cell culture remains unclear, it is clear that the syncytial plaque phenotype is favored under these conditions. Three independent lineages of Mixed viral populations became dominated by syncytial variants (**Figure 2**) under these experimental conditions, and the selective advantage of syncytia formation in Vero cell culture has been previously reported [44]. However, it is known that a syncytial phenotype can be caused by multiple independent genetic variants, and we can infer that each syncytial variant could have a different effect on viral fitness. We were able to identify two previously described syncytial variants in the Mixed viral population, both of which affect the gene encoding gB (UL27), at amino acids 817 (L to P) and 858 (R to H) [35–38] (**Figure 6**). Each of these variants steadily increased in frequency within the viral population over subsequent passages, but neither variant reached a majority of the population (and therefore became represented in the consensus genomes) until P8 (**Figure 6C**). As noted above, even when the visually observed phenotype indicated that the viral population was 88% syncytial (P7, **Figure 2A**), there was no genetic evidence of this at the level of the consensus genome. Intriguingly, while neither variant alone was sufficient to account for the entirety of this widespread syncytial phenotype, simply adding the frequency of each variant in the population neatly mirrored the frequency of syncytial plaques (**Figure 6C** and **2A**).

**Figure 6.**
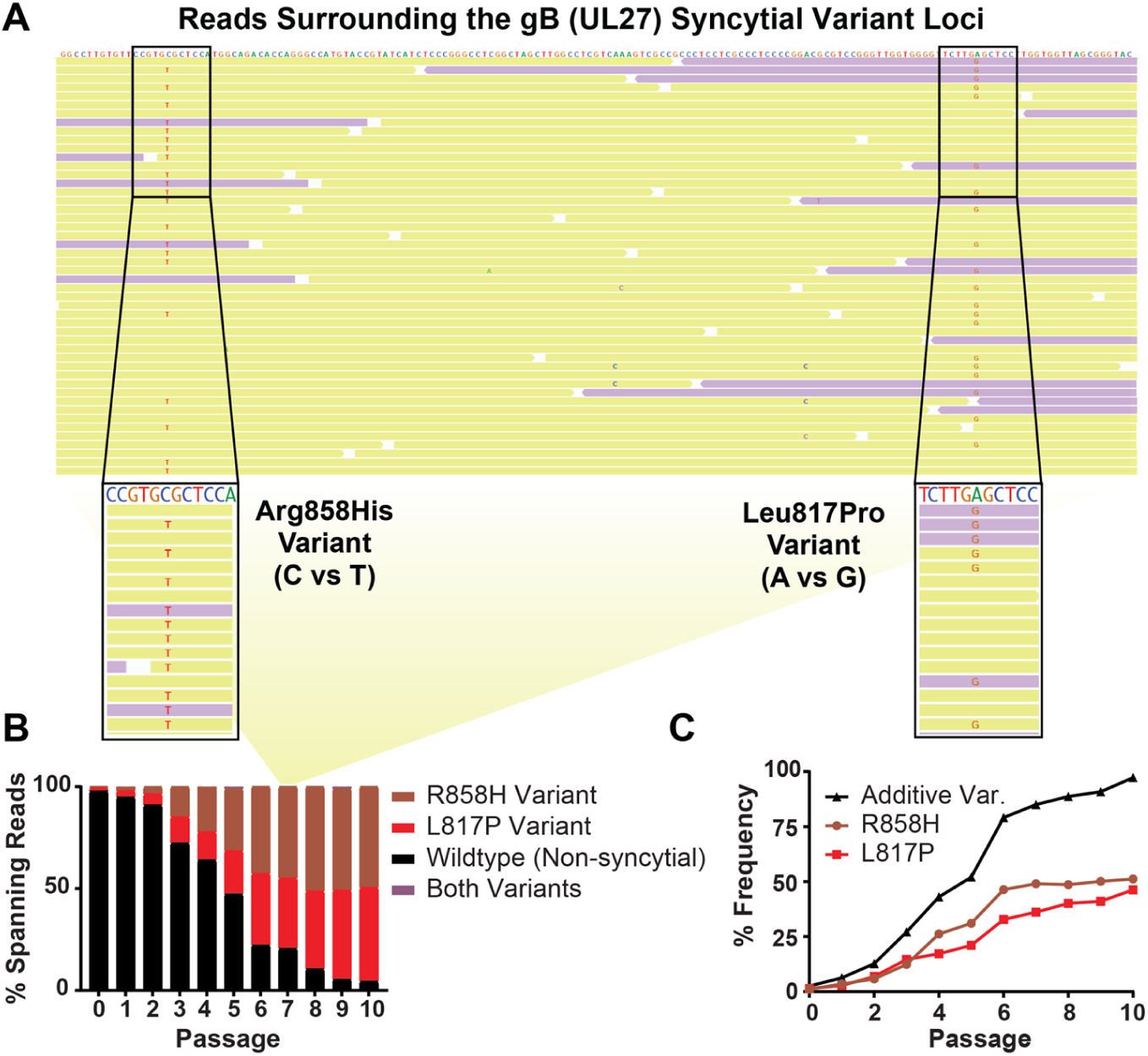
The Combination of Individual Variants in gB Explained the Syncytial Phenotype Observed in the Mixed Viral Population. The UL27 gene includes two syncytia-inducing minor variants that encode an Arginine to Histidine change at amino acid 858 or a Leucine to Proline change at amino acid 817. Sequencing reads that span the nucleotides encoding both sites were identified for each passage. **A)** Shown here is a random subset of the passage 7 (P7) sequencing reads surrounding this region, of which a total of 366 reads span both minor variants. The consensus genome is displayed at the top, with individual sequencing reads aligned below (yellow indicates forward-aligning sequencing reads, while purple indicates the reverse). Nucleotides matching the consensus genome are not shown, such that minor alleles that do not match the consensus sequence are the only ones highlighted. The relevant stretches of these sequencing reads are magnified, with any nucleotide that varies from the consensus genome noted (C-T for Arg858His or A-G for Leu817Pro). Note that the UL27 gene is on the reverse strand of the HSV-1 genome, so that the depicted nucleotides are in the reverse complement of what would be transcribed to produce the UL27 mRNA. **B)** Across all ten passages, the percent of sequencing reads spanning this region that contained one, both, or neither variant is summarized here. Data for “both” are included but are not visible on the graph, since only 6 sequence reads ever included both variants (three at P5, and three at P9). Yellow panel connects the subset of P7 sequence read data shown in **A)** to the relevant overall P7 percentage data graphed in **B). C)** The frequency of each variant was plotted alone, or as the sum of their frequency (Additive Variation (Var.)).

These data suggest that each of these syncytial variants in gB occurred and proliferated independently. Therefore, we sought to address whether or not these variants ever co-occurred on the same stretch of DNA. The proximity of these variants in the genome enabled us to identify a number of sequencing reads, each corresponding to one physical piece of DNA, that spanned both minor variant loci in gB (UL27) (**Figure 6A**). We detected each minor variant within this subset of reads at a frequency that was comparable to the overall frequency of the variant in the population (**Figure 6B-6C**). We found that each variant occurred independently. Across all reads spanning these loci in 10 passages, we found only 6 instances of these minor variants co-occurring in a single sequence read. We interpret these results as an indication that the syncytial phenotype, rather than any individual genetic variant, is being selected for under these conditions of Vero cell culture.

To test this hypothesis, we conducted a competition experiment in which two purified populations of HSV-1 were mixed in defined ratios and allowed to replicate under the same conditions as the original *in vitro* evolution experiment. We made use of a previously described syncytial clone of the F-Mixed population [27], which carries the R858H syncytial variant in gB, as well as the same F-Purified population as was used in **Figure 1**. These purified populations of uniformly syncytial or uniformly non-syncytial viruses were competed in defined ratios, where one or the other plaque phenotype was dominant at the start (50:1, 5:1, 1:5, and 1:50, Syncytial: Non-Syncytial). We also included one case of equal starting input (1:1, Syncytial: Non-Syncytial). As in the original *in vitro* evolution experiment, we found that viruses with a syncytial phenotype took over the viral population (**Figure 7**). The starting ratio influenced the rate of syncytial virus takeover, as shown by the slopes of each curve in **Figure 7**. However, all ratio mixtures were predominantly syncytial by P10.

**Figure 7.**
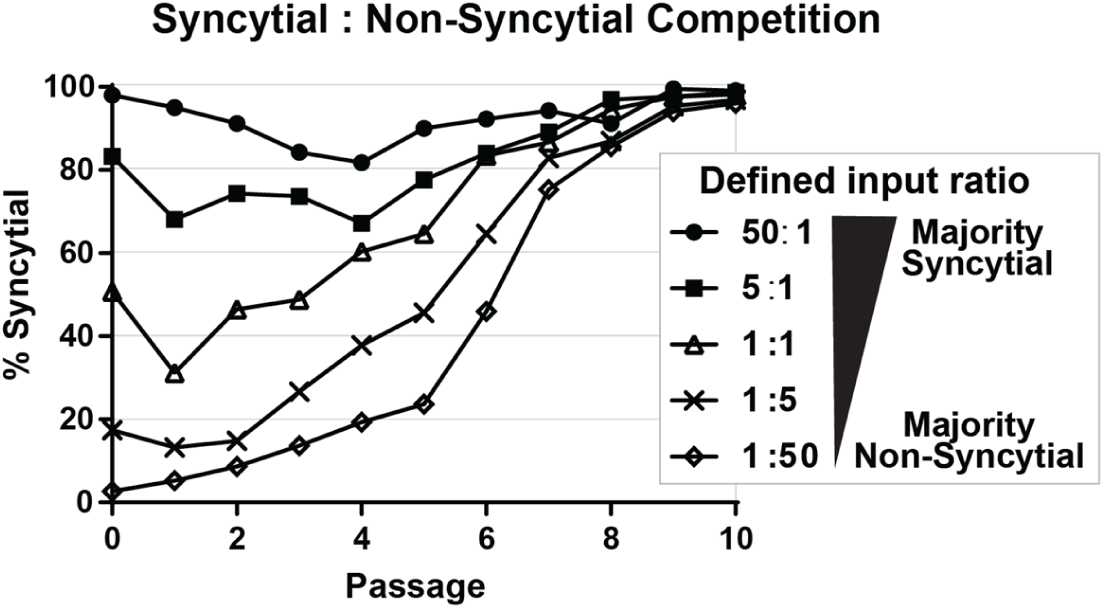
Defined Mixtures of Purified Syncytial and Non-Syncytial HSV-1 Populations Reveal a Universal Advantage of the Syncytial Phenotype during Sequential Passage in Vero cells. Purified populations of HSV-1 that displayed a uniformly syncytial or non-syncytial phenotype were mixed in defined ratios (as shown in legend) and allowed to compete during sequential passages of infection on Vero cells. At each passage, the plaque morphology of the resulting viral population was counted and plotted as the percentage of total plaques that were syncytial. MOI (0.01) and time to harvest (72 hpi) of each passage are the same as the initial *in vitro* evolution experiment described in **Figure 1**.

### Genetic Variation Outside of Syncytia Formation Facilitates Viral Fitness

Finally, we examined whether or not each evolved viral population was more fit for replication than the starting population. To do this, we compared the single-step and multiple-step growth kinetics for representative populations of the passaged viruses. We chose P0, P5, and P10 as representative of the Mixed viral population at the starting, midpoint, and endpoint of the *in vitro* evolution experiment. For the Purified viral population, we examined only the P0 and P10 populations, due to the limited genetic changes observed in that viral population. Under both single-step growth conditions (MOI=10), and multiple-step growth conditions (MOI = 0.01), we found that the passaged viral populations increased in titer to a greater degree than their original starting populations (**Figure 8**). The differences in replication between passaged and input viral populations were statistically significant (p < 0.05) at the final two time points in both growth curves (Two-way ANOVA), although the final titers of these stocks were overall quite similar (**Figure 8** and **Figure 2B**).

**Figure 8.**
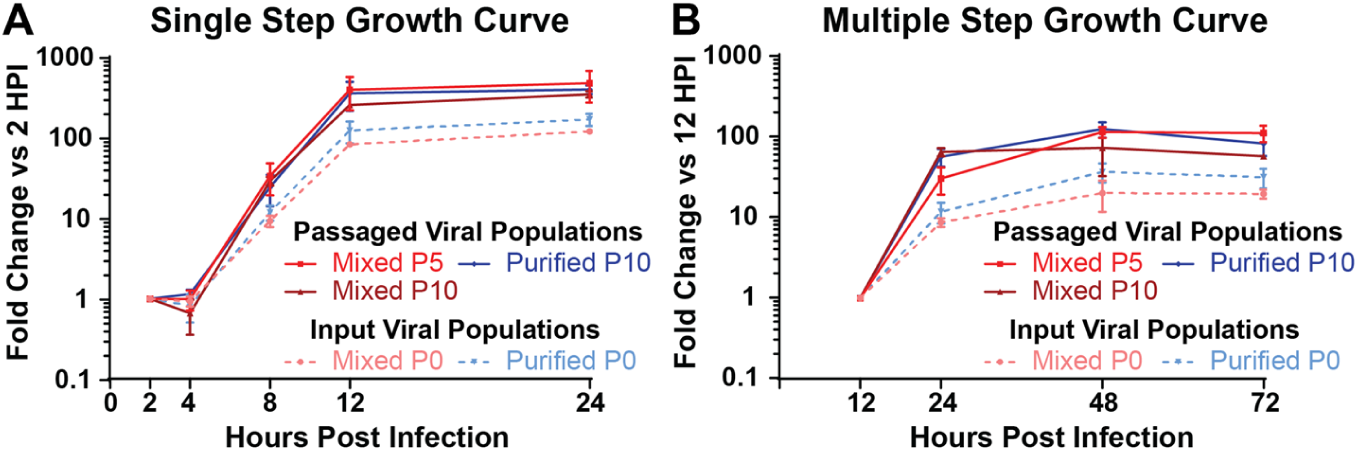
Passaged Virus Populations Replicate More than Input Viral Populations. Viral populations from the indicated passages of the Mixed or Purified stocks were used to infect Vero cells in **A)** a single step growth curve (MOI=10), or in **B)** a multiple step growth curve (MOI=0.01). Each assay was done in triplicate, with viral harvest and quantification by titering at the indicated timepoints. Data is plotted as titer compared to the first timepoint for each experiment (2 hours for **A**, 12 hours for **B**). Error bars indicate standard deviation. In both growth curves, comparisons of passaged viral populations versus the input viral populations were significantly different at the final two timepoints (Two-way ANOVA, p < 0.05).

## Discussion

In this study, we examined the evolution of HSV-1 during viral replication *in vitro*. We sequentially passaged either a mixed or a purified population of HSV-1 in Vero cell culture, and we quantified the outcome both in terms of observed phenotypes and in genomic changes over time. These approaches enabled us to monitor how the viral population changed from passage to passage, to detect potentially advantageous viral genotypes and/or phenotypes under these conditions, and to examine how a diverse versus a homogeneous pool of viral genomes impacted the subsequent genetic drift of each viral population. We found that the more diverse (“Mixed”) viral population was much more prone to change during passaging, with a syncytial plaque phenotype emerging as an advantageous trait under these Vero cell culture conditions. In addition to the genetic variants associated with syncytia formation, we observed the proliferation of a multitude of other genetic variants that may also confer a selective advantage in Vero cell culture conditions. Because Vero cell culture is the main methodology behind HSV-1 and HSV-2 propagation, both in experimental and diagnostic lab settings, these data provide key insights on the need to detect and understand viral evolution and selection *in vitro*. This is especially critical if the goal is to use laboratory observations to improve our understanding of viral population dynamics in actual human infections.

From these experiments, we were able to gain a number of useful insights for HSV-1, which may well extend to other herpesviruses and large DNA viruses. First, these experiments emphasized the contributions of standing variation and genetic diversity to HSV-1 evolution. While few would suggest that viruses with a large dsDNA genome do not evolve, the prevailing sentiment is that DNA viruses have a substantially slower rate of evolution than RNA viruses [5,45]. Thus, the concept of quasispecies or of viruses existing as populations with a mixture of genotypes has generally been limited to the RNA virus world. However, recent experiments that take advantage of modern deep sequencing technology have shown that dsDNA viruses possess substantial genetic diversity *in vivo* as well as in laboratory conditions [11,17,20,26,46–51]. The diversity of genomes in an RNA virus population has been linked to viral fitness [52,53], and there is no reason to expect a different result with DNA virus populations. Indeed, in these experiments, the genetically diverse (“Mixed”) viral population was able to adapt more readily to replication in Vero cells, while the more homogeneous (“Purified”) population appeared to be much more stable.

From a technical standpoint, these experiments highlighted the usefulness of deep sequencing for the analysis of viral populations. If we were to assume that the consensus genomes of each Mixed passage were indicative of the entire viral population, we could not have explained why there were more syncytial plaques observed over sequential passages, because the consensus genome did not change until passage 8 (P8). A focus solely on consensus-level genome analysis would also have missed the dynamics of the inactivating variants of UL13, not to mention the array of other genetic variants that fluctuated in the Mixed viral population (**Figure 5A, Table 1**). Limiting the genetic analysis to the consensus level, particularly when using deep sequencing approaches, eliminates a great deal of useful information about the viral population [14]. When performing similar evolution experiments to those performed here, but with different viruses, other researchers have found the most compelling evidence of evolution at the minority variant level [47,54–57].

Several of the specific minor variants observed in this study are ones that have been previously noted to occur spontaneously in HSV-1 and related herpesvirus strains during propagation *in vitro*. For instance, the occurrence of spontaneous syncytia-inducing mutations in the gene encoding gB (UL27) has been well-documented in the literature [35–38]. Likewise, the appearance of mutations that truncate the UL13 kinase, whether through an early stop codon or a frameshift, have been previously noted in multiple strain of HSV-1 by our lab and others [21,23,27] (**Figure 5B**). Despite prior observation in the literature, earlier works have not connected these variants to a fitness advantage in Vero cell culture per se. Future work is now warranted to investigate the potential positive selection of these and the other minor variants detected here that rise in frequency over sequential passages (e.g. **Figure 5A**). The rapid rise in frequency of UL13 variants observed in early passages of two clinical isolates (**Figure 5D**) suggests that the initial passages of new clinical isolates may serve as a useful screening approach to detect which of these minor variants are most impactful for viral fitness *in vitro* – i.e. those most frequently observed and/or rapidly selected in newly cultured clinical isolates. Intriguingly, both the UL13 and UL14 variants observed here were also detected in the initial culture passages of the genital HSV-1 isolate v29 [12].

These data lay the foundation for a number of subsequent explorations using similar approaches. First, as mentioned above, it is a goal of our lab and others to model the dynamics of HSV-1 infection that occur in human patients in a controlled setting of laboratory culture. Ideally, this controlled setting would minimize the genetic drift observed here in Vero cell culture. To minimize HSV-1 genetic drift, it may be useful to explore using alternative cell lines (e.g. species-matched or with intact interferon signaling, both of which are missing in Vero cells) or by using a different multiplicity of infection, or by changing the duration of infection. The *in vitro* evolution approach could also be used to investigate HSV-1’s response to specific selective pressures, such as exposure to antibodies, complement, or antiviral drugs. This type of *in vitro* evolution experiment has been performed with other viruses [47,54,55,58], though more frequently with RNA viruses rather than DNA viruses. While conventional reverse genetic approaches allow for detailed examination of the function of a specific gene product, forward genetic approaches such as *in vitro* evolution passaging experiments will allow us to study how the entirety of the viral genome responds to selective pressures. In addition, the unbiased approach of forward genetics enables the exploration of individual missense variants, as well as epistatic interactions of variants, and even copy number variants, as seen as prior studies [47,54,55,58]. These experiments may be especially fruitful in the case of large DNA viruses like HSV-1, where multiple genes have pleiotropic and/or unknown functions, or undefined networks of interactions.

## Materials and Methods

### Cells and Viruses

Vero (African Green Monkey Kidney) cells (ATCC, CCL-81) were maintained at 37C with 5% CO_2_. Cells were cultured in Dulbecco’s minimal essential medium (DMEM) supplemented with 10% Fetal Bovine Serum (FBS), penicillin-streptomycin (Life Technologies – Gibco) and L-Glutamine (Thermo Fischer Scientific-HyClone). HSV-1 strain F has been previously described [26,27]. In our previous description of mixed plaque morphologies in this stock of HSV-1 F, we referred to this stock as “F_Original_”, and here we refer to it as F-Mixed or Mixed throughout the paper. The Purified population used here is the same as the previously described F_Large_ [26]. For **Figure 7**, the purified Syncytial population (F-Syncytial) is the same as the previously described F_Syncytial_ [26]. The consensus genome sequence of the Purified (F_Large_) population is GenBank accession KM222724 and that of the purified Syncytial population (F_Syncytial_) is GenBank accession KM222725. Clinical isolates shown in **Figure 5** were collected at University of Washington from participants within one year of their first episode of genital HSV-1 infection. Isolates were collected from genital lesions, placed into viral transport media, and cultured once on Vero cells (Passage 0, P0) before transfer of samples to the Pennsylvania State University for further studies.

### Ethics Statement

The University of Washington Human Subjects Division reviewed and approved the collection of clinical isolates, and all participants provided written informed consent.

### *In Vitro* Evolution Experiments

Each viral population was used to infect a T-150 flask of Vero cells at an MOI of 0.01 in DMEM with 2% FBS, penicillin-streptomycin, and L-Glutamine. 72 hours post infection (hpi) virus was harvested by scraping, followed by three cycles of freezing and thawing. Each cycle of infection and harvest was considered a passage. The harvested virus was then titered on Vero cells using DMEM with 2% FBS, penicillin-streptomycin, L-Glutamine, and methylcellulose. At 72 hours post infection, cell monolayers were fixed and stained with methanol and methylene blue. Following a determination of titer, plaque morphologies were identified by counting at least 200 plaques per passage. This process then continued through each of 10 passages to create a “lineage”. For the Mixed population, three independent lineages were created.

### Other Viral Infections

For the competition experiments in **Figure 7**, Vero cells were infected at an MOI of 0.01, at the indicated ratios of F-Purified: F-Syncytial. For the growth curves in **Figure 8**, Vero cell monolayers were infected at a MOI of 10 (single-step) or 0.01 (multiple-step) and harvested by scraping at the indicated timepoints. Following three rounds of freezing and thawing, the resulting virus stock was titered on Vero cells as above. These experiments were performed in triplicate.

### Next-Generation Sequencing

Next-generation sequencing was performed as previously described [26]. Briefly, nucleocapsid DNA was prepared by a high (MOI=5) infection of Vero cells, followed by collection of cells and media. Nucleocapsids were isolated with Freon and pelleted through a glycerol gradient. Nucleocapsids were then lysed using SDS and Proteinase K, and viral DNA was extracted using phenol-chloroform. The viral nucleocapsid DNA was then used to prepare barcoded sequencing libraries, according to the Illumina TruSeq DNA Sample prep kit instructions. Libraries were quantified and assessed by Qubit (Invitrogen, CA), Bioanalyzer (Agilent), and library adapter qPCR (KAPA Biosystems). Illumina MiSeq paired-end sequencing (2 x 300 bp) was completed according to manufacturer’s recommendations, using 17 pM library concentration as input to the MiSeq. A consensus viral genome for each strain was assembled using a *de novo* viral genome assembly (VirGA) workflow [26]. Consensus genomes for each population were aligned and visualized using Geneious [59] (Biomatters). A multi-line fasta file of these consensus genomes, as well as the alignments used for **Figure 2**, are available in a ScholarSphere data repository at this DOI: https://doi.org/10.26207/t2vg-jd33.

### Minor Variant Analysis

Minor variants (MVs) within each genome were detected using VarScan 2.4.2 [60], in a similar manner as previously described [11,12,61]. We used parameters intended to eliminate sequencing-induced errors from the initial calling of MVs. Minimum variant allele frequencies ≥0.02; base call quality ≥20; read depth at the position ≥100; independent reads supporting minor allele ≥5. Minor variants with directional strand bias ≥90% were excluded. MVs that passed these filters were then annotated onto the genome using SnpEff [62] to determine their mutational effects. Finally, we used the alignment of the sequentially passaged consensus genomes to detect large gaps and/or insertions that were observed in single genomes, and to locally reassemble these suspected errors. Once curated, these genomes were re-analyzed for minor variants as before. See **Supplemental File 1** (Mixed viral populations) and **Supplemental File 2** (Purified viral populations) for full list of minor variants detected at each passage from P0-P10. High confidence minor variants (**Figure 5** and **Table 1**) were defined by the presence of a given variant in 3 or more consecutive passages in a lineage.

## Acknowledgements

The authors would also like to thank Nathan Arnett for his contributions to the passaging experiments, and the other members of the Szpara lab for their critical reading of the manuscript and figures. The authors acknowledge support from the Eberly College of Science and the Huck Institutes of the Life Sciences at Pennsylvania State University, as well as NIH grant support from R21 AI130676 and R01 AI132692 to MLS, and P01 AI030731 to CMJ. This project was funded, in part, under a grant with the Pennsylvania Department of Health using Commonwealth Universal Research Enhancement Program (CURE) funds (MLS).

## Supporting Information Legends

**Supplemental File 1. Minor Variants Identified in the Mixed Viral Population Passages:** The minor variants that were detected during deep sequencing of each passage of the HSV-1 F Mixed viral population are included as an Excel file. Substitutions and insertions/deletions are both listed, with one Excel sheet (or tab) for each passage from 0 (P0) through P10.

**Supplemental File 2. Minor Variants Identified in the Purified Viral Population Passages:** The minor variants that were detected during deep sequencing of each passage of the HSV-1 F Purified viral population are included as an Excel file. Substitutions and insertions/deletions are both listed, with one Excel sheet (or tab) for each passage from 0 (P0) through P10.

